# ALDH2 dysfunction accelerates ESCC pathogenesis

**DOI:** 10.1101/2023.04.03.534985

**Authors:** Samuel Flashner, Masataka Shimonosono, Norihiro Matsuura, Shinya Ohashi, Andres J. Klein-Szanto, J. Alan Diehl, Che-Hong Chen, Daria Mochly-Rosen, Hiroshi Nakagawa

**Affiliations:** Herbert Irving Comprehensive Cancer Center, Columbia University Irving Medical Center, Columbia University, New York, New York, USA; Department of Therapeutic Oncology, Graduate School of Medicine, Kyoto University, Shogoin, Sakyo-ku, Kyoto, Japan; Histopathology Facility, Fox Chase Cancer Center, Philadelphia, Pennsylvania, USA; Case Comprehensive Cancer Center, Department of Biochemistry, School of Medicine, Case Western Reserve University, Cleveland, Ohio, USA; Department of Chemical and Systems Biology, Stanford University School of Medicine, Stanford, CA, USA; Division of Digestive and Liver Diseases, Department of Medicine, Columbia University Irving Medical Center, Columbia University, New York, New York, USA

## Abstract

The alcohol metabolite acetaldehyde is a potent human carcinogen. Aldehyde dehydrogenase 2 (ALDH2) is the primary enzyme that detoxifies acetaldehyde in the mitochondria. Acetaldehyde accumulates and causes genotoxic stress in cells expressing the dysfunctional ALDH2^E487K^ mutant protein linked to *ALDH2*2*, the single nucleotide polymorphism highly prevalent amongst East Asians. Chronic alcohol users with heterozygous *ALDH2*2* display an increased risk for the development of esophageal squamous cell carcinoma (ESCC) and other alcohol-related cancers. However, how ALDH2 influences ESCC pathobiology is incompletely understood. Herein, we characterize how ESCC and preneoplastic cells respond to alcohol exposure using cell lines, three dimensional organoids, and xenograft models. We find that alcohol exposure results in increased organoid formation and tumor growth concurrent with increased reactive oxygen species (ROS), increased DNA damage, and the enrichment of putative cancer stem cells (CSCs) characterized by high CD44 expression. Pharmacological activation of ALDH2 function by Alda-1 inhibits this phenotype, indicating that acetaldehyde is the primary driver of these changes. ALDH2 dysfunction also affects response to a commonly used chemotherapy for the treatment of ESCC. We find that Aldh2 dysfunction facilitated enrichment of CSCs following cisplatin-induced cell death and oxidative stress in murine organoids. Together, these data provide evidence that alcohol exposure, results in more aggressive tumors through enrichment of CSCs, which is augmented by ALDH2 dysfunction.

## INTRODUCTION

Alcohol consumption is a leading risk factor for esophageal squamous cell carcinoma (ESCC), which accounted for ∼500,000 deaths worldwide in 2020 [1]. ESCCs harbor dismal five-year survival rates due to late diagnosis, metastasis, and therapy resistance [2]. There is an urgent need to characterize the factors that influence the pathobiology of this disease. Alcohol consumption exposes the esophageal mucosal surface to high concentrations of ethanol (EtOH) and acetaldehyde, a chief metabolite of EtOH [3,4]. After alcohol drinking, acetaldehyde levels in saliva are ten-times higher than in blood [5]. Because ESCCs are commonly diagnosed late and in individuals who chronically consume alcohol, understanding the effects of alcohol exposure on preneoplastic lesions as well as established tumors will illuminate salient features of a deadly carcinogenic process.

The tumor promoting effects of alcohol are thought to be derived from its metabolic intermediates, including acetaldehyde. Certain alcoholic beverages consumed in the high incidence areas for esophageal cancer contain >25% EtOH and >1 mM acetaldehyde [5,6]. Acetaldehyde is a highly reactive compound that can damage both mitochondria and DNA [7– 9]. Acetaldehyde is detoxified by the mitochondrial enzyme acetaldehyde dehydrogenase 2 (ALDH2) that catalyzes oxidation of acetaldehyde into acetic acid [10]. The ALDH2 rs671 (ALDH2*2) is a single nucleotide polymorphic mutation carried by 35-45% of East Asians and is among the most common ALDH2 polymorphisms worldwide. The dominant negative mutant ALDH2*2 allele encodes for the E487K change in the ALDH2 protein, which is associated with a dramatic decrease in enzymatic activity compared with wild type ALDH2*1 [11]. Interestingly, heavy drinkers who are also heterozygous carriers of the ALDH2*2 allele have a ∼14-fold increased risk for ESCC as well as worse prognosis compared to those who develop the disease without ALDH2*1 [12–14]. Despite the severe burden of this disease subtype, how acetaldehyde accumulation promotes worse prognosis in ESCC heterozygous ALDH2*2 carriers is unclear.

One key determinant driving ESCC pathobiology is the presence of putative cancer stem cells (CSCs) characterized by high expression of the cell surface glycoprotein, CD44 (CD44H) [15]. CD44H cells are therapy resistant, highly proliferative, metastatic, and capable of initiating tumors [16–21]. We have recently demonstrated that alcohol exposure promotes enrichment of CD44H cells that tolerate excessive levels of alcohol-derived ROS through increased autophagy [22]. How ALDH2 status influences alcohol-driven CSC enrichment has not been studied. We hypothesize that ALDH2 dysfunction underlies the observed CSC enrichment.

To address this hypothesis, we leveraged our recently developed three dimensional (3D) esophageal organoid platform. 3D organoids are formed from single cells and capture the functional, morphological, and molecular features of the original tissue [16,23–28]. Key to this study, 3D organoids harbor a distinguishable population of CD44H cells that more readily form tumors as well as organoids compared with their counterparts of low CD44 expression (CD44L) [16,23–25].

Herein, we examine the pathogenic consequences of ALDH2 dysfunction during EtOH-induced ESCC carcinogenesis. We leverage 3D organoids derived from isogenic mice harboring either wild type ALDH2 or heterozygous ALDH2*2. In parallel, we utilize human ALDH2*2 ESCC 3D organoids and xenograft tumor models in the presence or absence of the ALDH2 activator Alda-1 [29]. Using these tools, we demonstrate that alcohol exposure promotes CSC enrichment and tumor growth in an ALDH2*2-dependent manner. These deleterious effects are inhibited by restoring ALDH2 function by the pharmacological ALDH2 activator, Alda-1 [29]. Moreover, we show that alcohol exposure generates ROS and DNA damage to enrich CSCs. We further demonstrate that the *Aldh2*2* polymorphism modulates response to a commonly used chemotherapy drug for the treatment of ESCC, cisplatin. Together, our data unravel a key axis driving the malignancy of a common and deadly disease.

## RESULTS

### Heterozygous ALDH2*2 potentiates ethanol-induced CSC enrichment

To evaluate how the Aldh2*2 polymorphism modulates ESCC pathobiology, we first established an isogenic model of malignant transformation. We treated Aldh2*1/Aldh2*1 homozygous wild-type (WT) mice and Aldh2*1/Aldh2*2 heterozygous mutant (HET) mice with the potent carcinogen, 4-Nitroquinoline 1-oxide (4-NQO), known to drive ESCC transformation in mice [16]. Following 4-NQO treatment, we collected esophagi from both cohorts of mice and evaluated the histological progression of ESCC. We observed no statistically significant difference in ESCC progression in Aldh2*2 HET compared with Aldh2 WT mice (Supplemental Figure 1B). We then used the remaining tissue to generate 3D organoids which retain the morphological features of their tissue of origin (Supplemental Figure 1B). We observed a slight increase in organoids harboring high grade dysplasia in the Aldh2*2 HET condition compared with Aldh2 WT organoids (Supplemental Figure 1C). Together, we established a tractable organoid model of ESCC pathogenesis in the context of Aldh2 dysfunction.

We next leveraged this 3D organoid system to determine how the Aldh2*2 polymorphism influences alcohol-induced pathogenesis. To capture the salient changes that occur during ESCC initiation, we used Aldh2*2 HET or Aldh2 WT organoids with histological features consistent with the ESCC precursor lesion intraepithelial neoplasia (IEN). Using this model, we first determined how the Aldh2*2 polymorphism may alter secondary organoid formation (OFR), a surrogate readout of tumor initiating capability [22]. To do so, we exposed Aldh2 WT and Aldh2*2 HET to 1% ethanol (EtOH) for 96 hours, dissociated organoids into single cells, passaged these cells and measured their ability to form organoids. We found that EtOH induced a nearly three-fold increase in OFR in Aldh2*2 HET organoids compared with a modest ∼1.2 fold change in Aldh2 WT organoids (Figure 1A). We next considered whether these changes in OFR were due to the enrichment of the CD44H putative CSC population within each organoid. To address this possibility, we treated organoids with EtOH as before, collected single cells, and performed flow cytometry for CD44H content. Consistent with our OFR results, we observed a nearly four-fold increase in CD44H cells in the Aldh2*2 HET organoids following EtOH exposure, compared with a two-fold increase in the Aldh2 WT organoids (Figure 1B). Together, these results indicate that EtOH exposure accelerates ESCC transformation in the context of Aldh2 dysfunction.

**Figure 1:**
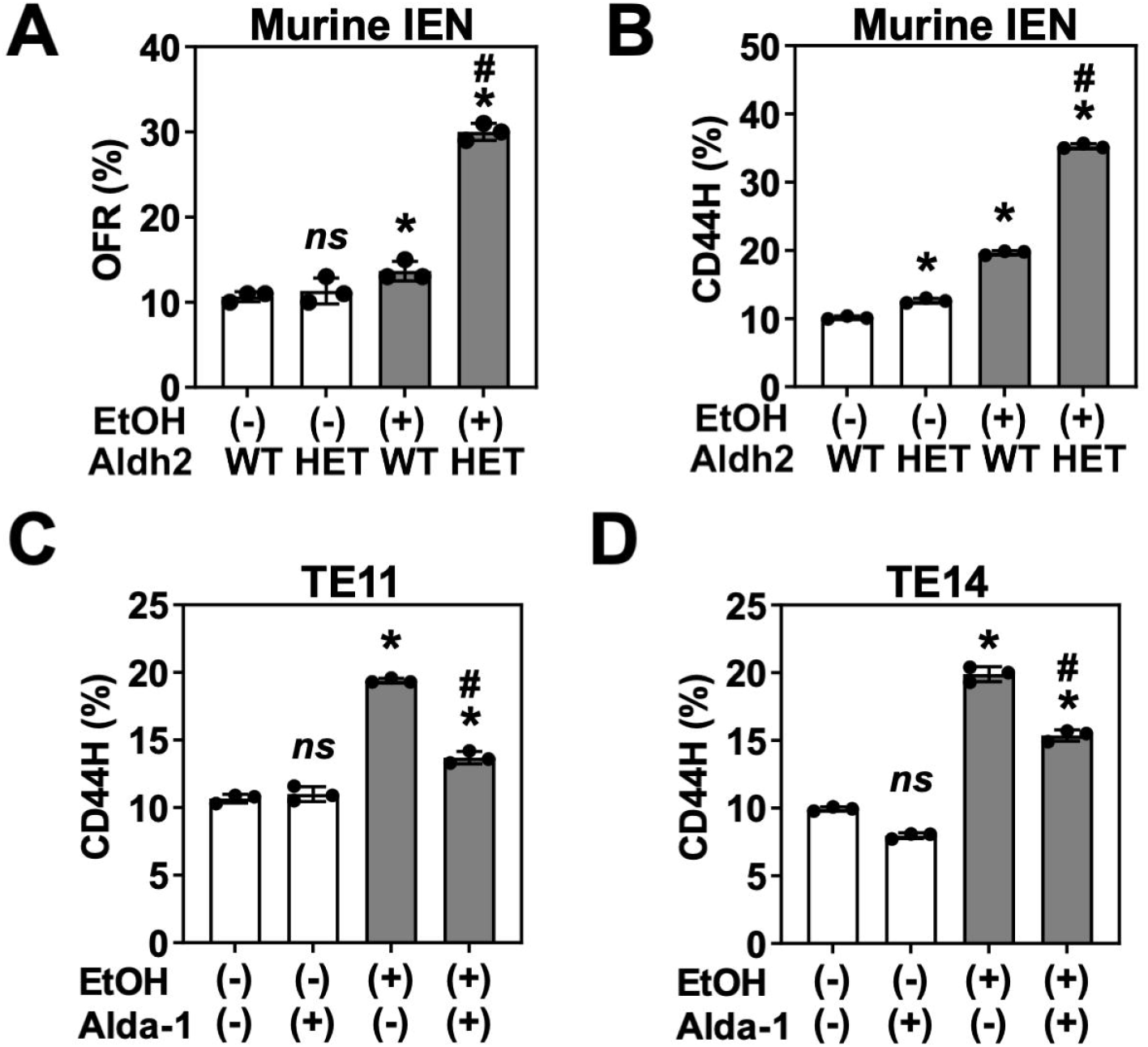
Heterozygous ALDH2*2 potentiates ethanol-induced CSC enrichment. Murine IEN organoids from the indicated genotype were treated with 1% EtOH from day 7 to day 11 of culture. Organoids were collected, dissociated into single cells, and then assessed for (**A**) secondary organoid formation by brightfield microscopy, (**B**) CD44H cell content by flow cytometry (FCM). (**C-D**) Organoids from two human ESCC lines heterozygous for ALDH2*2 (TE11 or TE14) were treated with 1% EtOH +/-20 µM of the ALDH2 activator Alda-1 from day 7 to day 11 of culture, collected, dissociated into single cells, and then subject to FCM for detection of CD44H cell content. ns = not significant.^*^p <0.05 relative to untreated control. ^#^p<0.05 relative to EtOH-treated ALDH2 WT control (**A-B**) or EtOH-treated, Alda-1-untreated cells (**E-F**). Circles represent technical replicates (n = 3).

We next addressed whether our findings translated to established human ESCC tumors. We used TE11 and TE14 cells, which are two independent ESCC cell lines heterozygous for the ALDH2*2 allele. To isolate the effect of ALDH2 dysfunction using these models, we restored ALDH2 function with the pharmacological activator Alda-1 [29]. We treated organoids made with TE11 and TE14 cells with 1% EtOH for 96 hours in the presence or absence of ALDH2 activator Alda-1. We then dissociated the organoids into single cells and performed flow cytometry for CD44. Consistent with our results in murine IEN organoids, we found that short-term EtOH exposure resulted in an increased burden of CD44H CSCs (Figure 1C and D). Alda-1 partially inhibited CD44H enrichment, indicating that ALDH2 dysfunction potentiates the tumor-promoting effects of EtOH on human ESCC pathobiology (Figure 1C and D).

### Ethanol treatment promotes CSC enrichment by promoting non-CSC apoptosis in the context of Aldh2 dysfunction

We next evaluated the mechanisms underlying CSC enrichment. We had previously demonstrated that EtOH promotes apoptosis in CD44 low expressing cells (CD44L), but not CD44H cells [22]. To determine if such a mechanism was driving CD44H cell enrichment in this model, we first confirmed that EtOH exposure results in increased apoptosis in Aldh2*2 HET cells. We grew Aldh2 WT and Aldh2*2 HET cells in monolayer culture, treated with 2% EtOH for 72 hours, collected single cells, and performed flow cytometry for Annexin V and Propidium Iodide (PI). We observed a significant increase in Annexin V positive/PI negative early apoptotic cells in the Aldh2*2 HET organoids exposed to EtOH compared with Aldh2 WT organoids (Figure 2A – 2B). To confirm that this change was driving the observed CSC enrichment (Figure 1B), we repeated this experiment and measured early apoptosis in CD44H and CD44L cells. We found that EtOH exposure resulted in a significant increase in early apoptotic CD44L cells compared with CD44H cells (Figure 2C). This effect was most pronounced in Aldh2*2 HET cells. Together, these data indicate EtOH exposure promotes CSC enrichment in the context of Aldh2 dysfunction by promoting apoptosis in non-CSC cell populations.

**Figure 2:**
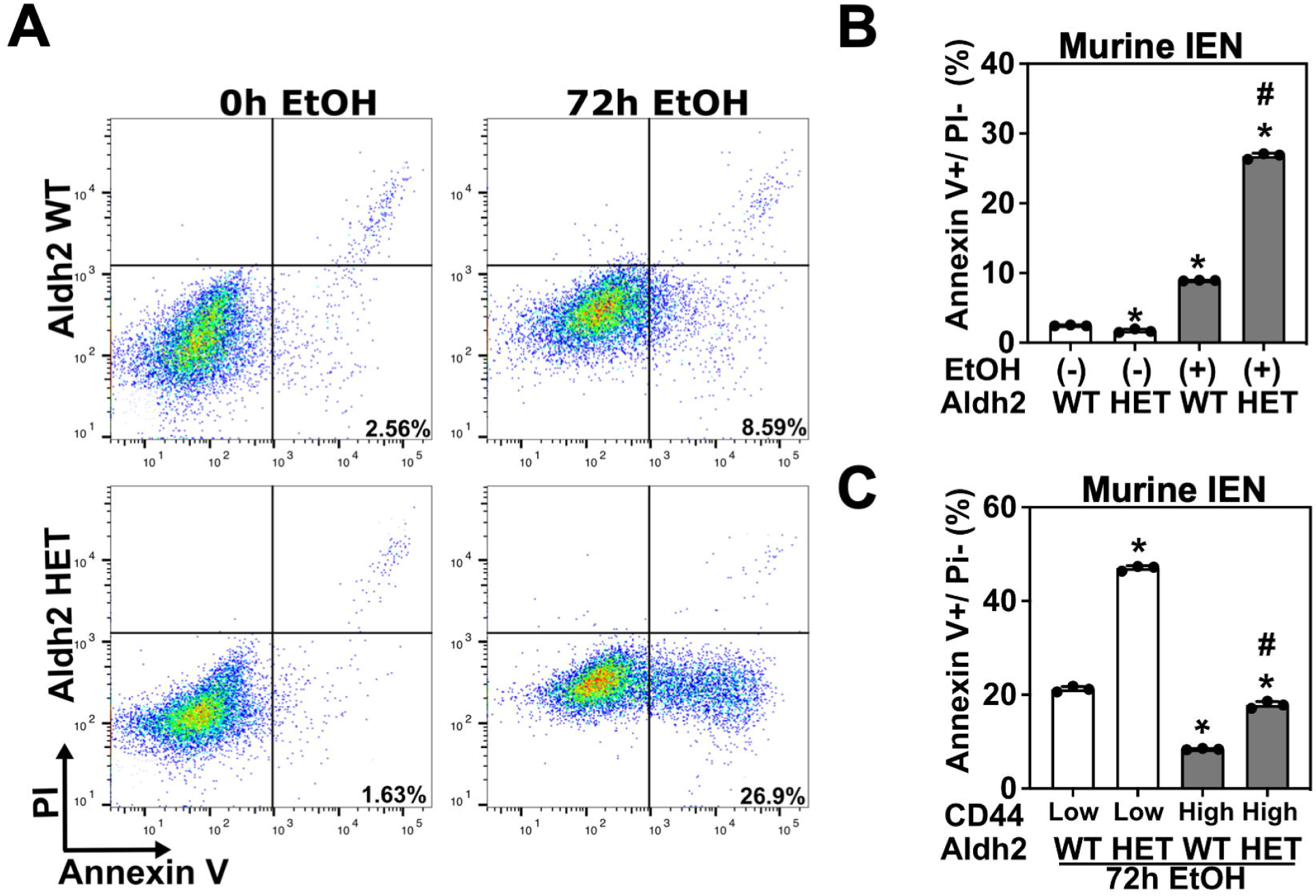
Alcohol promotes CSC enrichment by promoting non-CSC apoptosis in the context of Aldh2 dysfunction. Cells from the indicated genotype were grown in 2D, treated with 2% EtOH for 72 hours, collected, and assessed for apoptosis by FCM for Annexin V/PI. (**A**) representative dot plot of Annexin V and PI staining, (**B**) quantification of (**A**). (**C**) Apoptosis was measured in CD44H and CD44L cells by FCM. *p<0.05 compared with WT control (**B**) or with WT CD44L cells (**C**) #p<0.05 compared with HET control (**A**) or with HET CD44L cells (**C**). Circles represent

### Heterozygous ALDH2*2 potentiates ethanol-derived ROS accumulation and DNA damage

We next considered how alcohol may promote CSC enrichment in the context of Aldh2 dysfunction. We have previously demonstrated that the accumulation of reactive oxygen species (ROS) drives CD44H CSC enrichment [17,22]. We therefore assessed whether ROS accumulates in Aldh2*2 HET organoids. We treated organoids with 1% EtOH for 96 hours, dissociated organoids into single cells, and then measured mitochondrial ROS using the mitochondrial super oxide assay. We found that EtOH exposure results in a significant increase in mitochondrial ROS in Aldh2*2 HET organoids compared with EtOH treated Aldh2 WT organoids (Figure 3A). Furthermore, untreated Aldh2*2 HET organoids exhibit a modest increase in mitochondrial ROS compared with Aldh2 WT controls, highlighting the potentially deleterious effects of Aldh2 dysfunction independent of EtOH exposure. Together, these results indicate that Aldh2 dysfunction augments ROS production following EtOH exposure.

**Figure 3:**
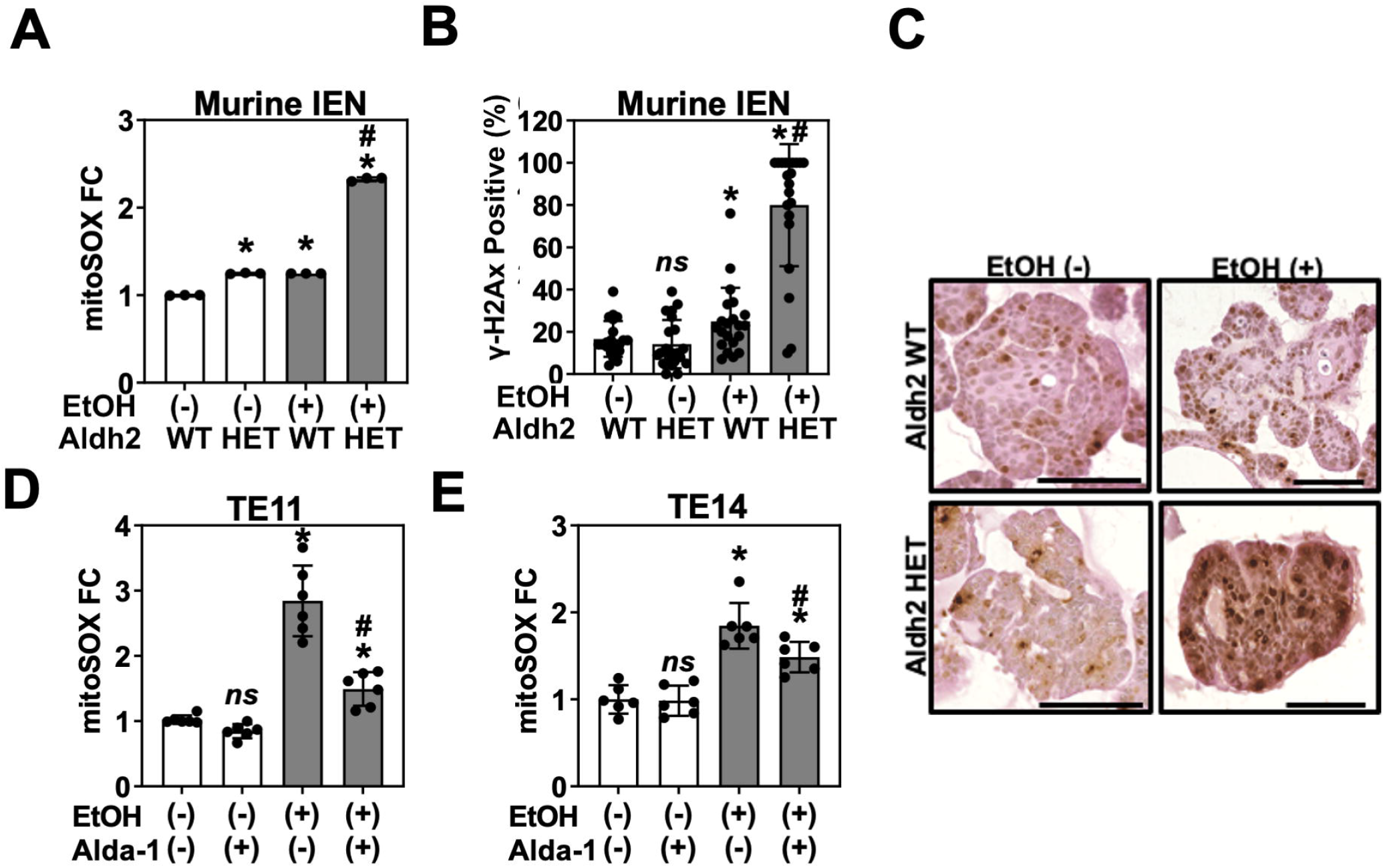
Heterozygous ALDH2*2 potentiates ethanol-derived ROS accumulation and DNA damage. Murine IEN organoids from the indicated genotype were treated with 1% EtOH from day 7 to day 11 of culture. Organoids were collected and (**A**) dissociated into single cells to assess mitochondrial ROS via the mitoSOX assay or (**B, C**) stained for yH2Ax and scored for marker positivity by a board-certified pathologist. TE11 or TE14 cells were grown in monolayer culture and then treated with 1% EtOH +/-20 µM of the ALDH2 activator Alda-1 for 24 hours. Mitochondrial ROS levels were assessed via fluorescence microscopy. ns = not significant.^*^p <0.05 relative to untreated control. ^#^p<0.05 relative to EtOH-treated ALDH2 WT control (**A-B**), EtOH-treated, Alda-1-untreated cells (**D** -**E**). Circles represent technical replicates (n ≥ 3). Scale = 100 µm.

Mitochondrial ROS exert their deleterious effects in part through oxidative DNA damage. We have previously demonstrated that EtOH exposure results in increased DNA damage in the esophageal epithelia of Aldh2^-/-^ mice [30]. To confirm that the increase in ROS has a functional effect on genome integrity in IEN organoids, we performed immunohistochemistry for the DNA double strand break marker y-H2Ax in IEN organoids treated with 1% EtOH for 96 hours. Consistent with the results of our mitochondrial superoxide assay, we found that EtOH exposure results in a significant increase in the percentage of y-H2Ax positive cells in the context of Aldh2 deficiency, with ∼80% of cells positive for y-H2Ax in Aldh2*2 HET mice treated with EtOH compared with <20% rate in untreated controls (Figure 3B, C). Further, we observed a modest significant increase in y-H2Ax positivity in Aldh2*1 cells, indicating that EtOH exposure influences ROS and DNA damage in the context of Aldh2 competency, albeit to a lesser extent than in the Aldh2*2 HET organoids. Together, these data support our hypothesis that ALDH2 limits the accumulation of CSC-enriching ROS following EtOH exposure.

We next assessed whether EtOH exposure results in a similar level of ROS induction in human ESCC cell lines heterozygous for ALDH2*2. We treated TE11 and TE14 cell lines grown in monolayer with 1% EtOH and 20 µM Alda-1 for 24 hours and measured mitochondrial ROS via the mitochondrial superoxide assay. We observed a significant increase in ROS in these ALDH2-dysfunctional cell lines following EtOH exposure, which was reversed following ALDH2 activation by Alda-1 (Figure 3D). These results are consistent with our above-described findings in murine IEN organoids. Together, these data corroborate our findings in murine IEN organoids that ethanol exposure promotes CSC enrichment and ROS accumulation during ESCC pathogenesis.

### Heterozygous ALDH2*2 accelerates ethanol-induced ESCC pathogenesis

We next addressed the translational significance of our findings. How the ALDH2*2 polymorphism influences the pathobiology of established tumors following alcohol consumption is undefined. We previously demonstrated that alcohol consumption promotes the growth of xenograft tumors in mice bearing the ESCC cell line TE14 [22]. We therefore evaluated whether ALDH2 dysfunction is underlying this phenomenon by subcutaneously transplanting TE14 into the flanks of athymic nu/nu mice. We treated mice with drinking water containing 10% EtOH and 20 mg/kg/day Alda-1 for six weeks. We observed a statistically significant increase in tumor size in the EtOH-treated mice compared with vehicle controls (Figure 4A). Alda-1 treatment inhibited the growth-promoting effect of EtOH, indicating that ALDH2 dysfunction is required for this process (Figure 4A). We next harvested these tumors and performed flow cytometric analysis of their CD44H cell content. We observed a statistically significant increase in the percentage of CD44H cells in ethanol exposed mice, which was inhibited following treatment with Alda-1 (Figure 4B). These data indicate that ALDH2*dysfunction supports ethanol-dependent CSC enrichment in vivo. Together, our work demonstrates that ethanol exposure promotes tumor growth and C44H CSC enrichment in the context of ALDH2*2 dysfunction.

**Figure 4:**
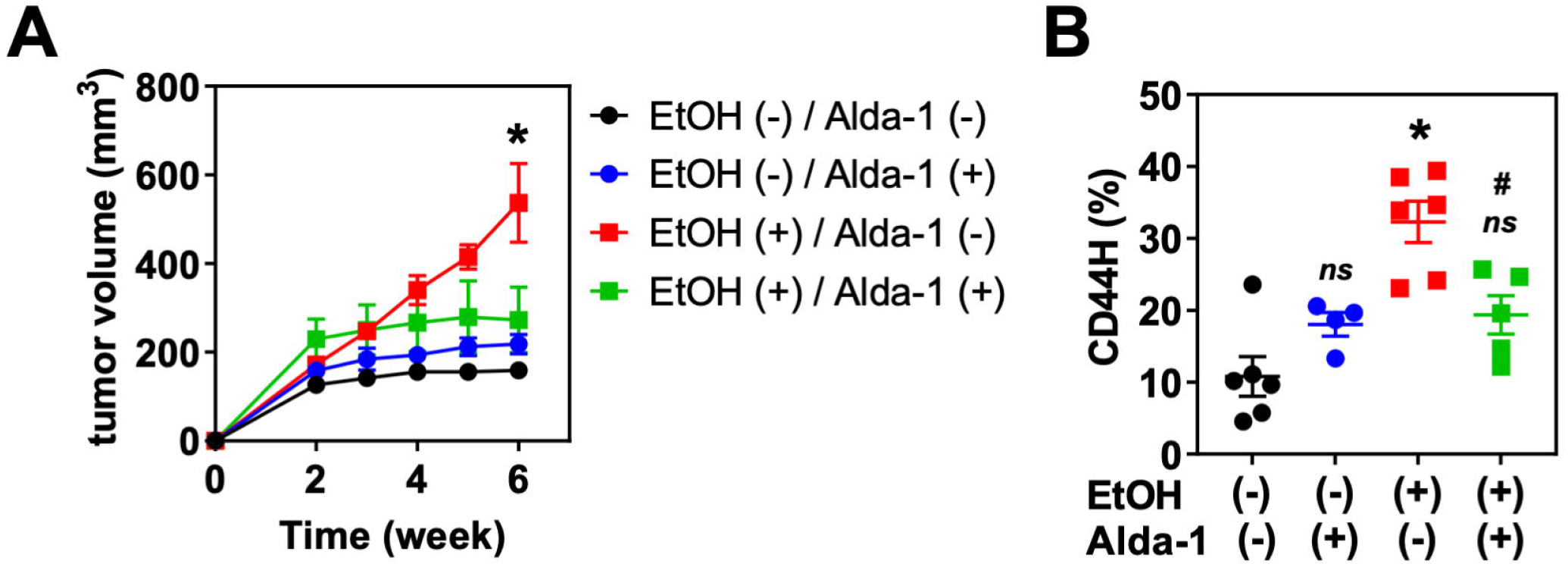
Heterozygous ALDH2*2 accelerates ethanol-induced ESCC pathogenesis. TE14-RFP cells were injected into the flanks of nu/nu mice. 10% EtOH was included in the drinking water ad libitum from week 2 to week 6 following transplantation. During the EtOH treatment period, Alda-1 (**20 mg/kg**) or vehicle control) was intraperitoneally injected twice daily. (**A**) Tumor volume was measured weekly. (**B**) At week 6, tumors were dissociated into single cells and analyzed by FCM for CD44. ns = not significant. ^*^p<0.05 compared with untreated control. ^#^p<0.05 compared with EtOH (+) / Alda-1 (-) tumors. Circles represent technical replicates (n ≥ 4 for both (**A**) and (**B**)).

### Heterozygous ALDH2*2 potentiates cisplatin-induced CSC enrichment

We next assessed how Aldh2 status may influence response to other genotoxic agents. Cisplatin is a commonly used chemotherapy for the treatment of ESCC. Since both acetaldehyde and cisplatin induce DNA interstrand crosslinks, a common precursor lesion for DNA double strand breaks, we hypothesized that Aldh2 dysfunction will promote CSC enrichment and ROS accumulation in response to cisplatin. To address this hypothesis, we performed a dose response with cisplatin in Aldh2 WT and Aldh2*2 HET organoids for 72 hours. We collected organoids, dissociated into single cells, and measured cell viability, CD44 levels, and mitochondrial ROS by flow cytometry. We found that the Aldh2*2 HET organoids were more sensitive to cisplatin, with ∼90% of cells dying following treatment with 80 µM cisplatin compared with ∼12% of Aldh2 WT cells (Figure 5A). We next evaluated whether cisplatin alters ESCC pathogenesis in a similar mechanism as ethanol exposure (Figures 1-4). Indeed, we observed a cisplatin dose-dependent increase in ROS in Aldh2*2 HET organoids compared to both untreated and Aldh2 WT controls (Figure 5B). Finally, we evaluated whether Aldh2 dysfunction promoted CD44 cell enrichment. We observed a significant increase in CD44H cell levels in Aldh2*2 HET organoids following exposure to any dose of cisplatin (Figure 5C). A modest increase in CD44 levels also occurred in Aldh2 WT organoids, corroborating the previous reports that cisplatin results in increased CSCs [31–33]. Together, these data indicate that Aldh2 status influences response to commonly used chemotherapy for the treatment of ESCC.

**Figure 5:**
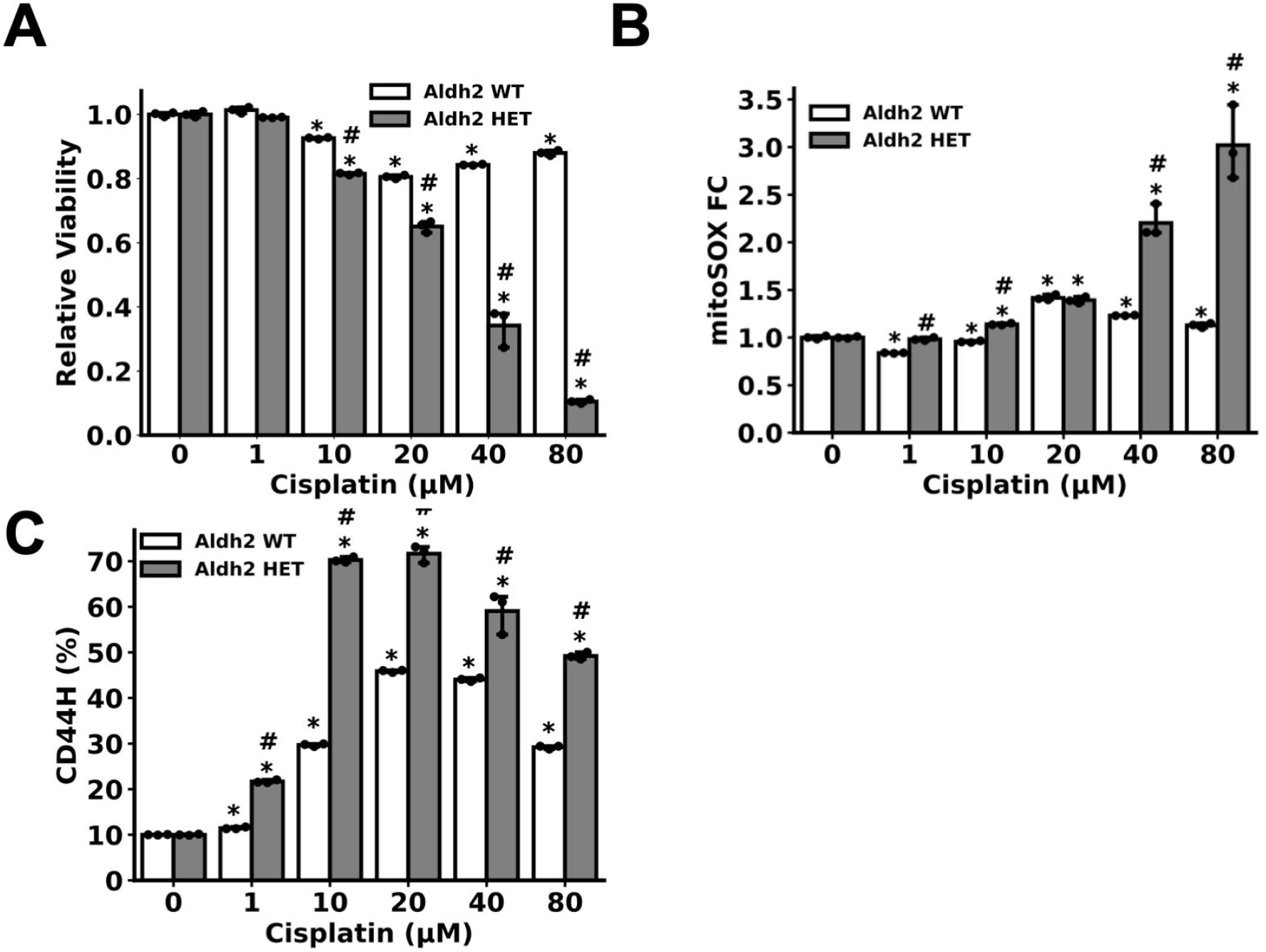
Heterozygous ALDH2*2 potentiates cisplatin-induced CSC enrichment. Murine IEN organoids from the indicated genotype were treated with the indicated concentration of cisplatin for 72 hours. Organoids were collected, dissociated into single cells, and subjected to FCM to assess for (**A**) viability by measuring DAPI exclusion, (**B**) mitochondrial ROS through the mitoSOX assay and (**C**) CD44H cell content. ^*^p<0.05 relative to untreated control. ^#^p<0.05 relative to Aldh2 WT sample treated with identical treatment of cisplatin. Circles = technical replicates (n = 3).

## DISCUSSION

Alcohol-related ESCC is a common and deadly disease, particularly in patients heterozygous for ALDH2*2 [14,34]. Understanding how alcohol influences ESCC pathogenesis in the context of ALDH2 dysfunction is therefore paramount to identifying the factors underlying the poor prognosis of these patients and identifying new therapeutic strategies to ameliorate the burden of this disease. Here, we demonstrate that ALDH2 dysfunction potentiates alcohol-induced tumor growth, ROS accumulation, DNA damage, and the enrichment of CSCs. Moreover, we find that ALDH2 dysfunction increases the response to a commonly used chemotherapy, cisplatin. Together, these data indicate that the common ALDH2*2 single nucleotide polymorphism accelerates ethanol-induced pathogenesis and influences response to cisplatin therapy.

Our data further the understanding of how ALDH2*2 polymorphism promotes malignant transformation following ethanol exposure. The preponderance of studies to date have focused on how ethanol exposure promotes malignant transformation by inducing DNA damage and genome instability [35–38]. In this model, ethanol-induced activation of oncogenes or inactivation of tumor suppressors drives ESCC tumorigenesis. We augment this understanding of ESCC pathogenesis by demonstrating that brief exposure to ethanol (24 hours in culture or 6 weeks in xenograft models) induces a dramatic change in the composition of a tumor or premalignant lesion. We demonstrate that ethanol enriches tumor-initiating CSCs in ESCC organoids and xenograft tumors, as well as their histologic precursor lesion IEN (Figure 1, Figure 4). We confirm that cells positive for the CSC marker CD44H do recapitulate the hallmark features of CSCs. In this study, organoid/cell populations with increased CD44H content form larger xenograft tumors (Figure 4), form organoids at a higher rate (Figure 1) and are enriched following treatment with chemotherapeutic agents (Figure 5) than populations with low levels of CD44H cells. These findings are all consistent with known features of CSCs, which can form more aggressive and faster growing tumors which are resistant to a variety of drugs.

One key finding of the present work is that alcohol-induced CSC enrichment occurs during IEN. While CD44H cells are appreciated to emerge during the premalignant stages of ESCC pathogenesis [39], this work is the first to demonstrate that this enrichment occurs in response to ethanol exposure in the context of ALDH2 dysfunction. Furthermore, we find that treatment of IEN organoids with chemotherapy enriches for CSCs (Figure 5), which tolerate higher levels of cisplatin in other cancer contexts and therefore drive resistance to the drug [31–33]. The presence of this highly aggressive and putatively drug-resistant subset of cells in a pre-malignant lesion (Figure 1B) highlights the need for earlier detection of ESCC pathogenesis. In addition to ethanol-induced CSC enrichment, we observe a modest but significant increase in CD44H cells in untreated Aldh2*2 HET IEN organoids compared with Aldh2 WT controls (Figure 1B). These data indicate that Aldh2 status may influence ESCC pathogenesis independent of ethanol exposure and is the focus of ongoing work in the lab. Ultimately, the work herein augments our current understand of how ethanol and ALDH2*2 influence ESCC pathogenesis. We demonstrate that these factors interact to alter the fundamental makeup of these tumors by supporting the enrichment of CSCs associated with poor patient prognosis and response to chemotherapy.

We hypothesize that the ethanol- and cisplatin-induced ROS accumulation and genotoxic stress represent selective pressures that enrich existing CSCs in the tumor or IEN microenvironments. We have previously demonstrated that ROS or genotoxic stress mediates CD44H enrichment in a variety of cancer contexts [16,18,22]. Moreover, cisplatin is known cause CSC enrichment in other cancer contexts [31–33] as well as ROS accumulation in the context of Aldh2 dysfunction [40]. Supporting our hypothesis, ethanol exposure results in apoptosis in CD44L, but not CD44H cells (Figure 2C-D). How CSCs survive elevated ROS/genotoxic stress is unclear. We have previously demonstrated that CD44H cells activate autophagy to clear ROS that arises from various sources, including alcohol [17,22]. In other contexts, we have demonstrated that CSCs survived otherwise cytotoxic levels of ROS by upregulating antioxidants such as SOD2 [18]. Precisely how CSCs survive elevated genotoxic stress is unclear and represents a limitation of this study.

That ethanol exposure results in increased ROS in the context of ALDH2 dysfunction is well documented. ALDH2*2 carriers cannot efficiently metabolize acetaldehyde, allowing its accumulation following ethanol exposure. Acetaldehyde can directly induce ROS [41]. However, how Aldh2 dysfunction results in increased ROS accumulation following cisplatin exposure is not known. One potential explanation is that ALDH2*2 carriers harbor basal levels of mitochondrial damage arising from the inability to metabolize endogenous aldehydes. This increased basal level of ROS may render cells susceptible to increased cisplatin-mediated ROS. Our data support this hypothesis. We observe a modest but significant increase in mitochondrial ROS in untreated Aldh2*2 HET organoids compared with Aldh2 WT organoids (Figure 3A). This increase may further explain the elevated CD44H cell content that we observed in untreated Aldh2*2 HET organoids compared with Aldh2*1 WT organoids (Figure 1B), although more work is needed to strengthen this claim.

These findings may translate beyond ESCC. ESCC and head and neck squamous cell carcinomas (HNSCC) may develop concurrently or independently in chronic alcohol users heterozygous for ALDH2*2 [42]. Furthermore, HNSCC ALDH2*2 HET patients who consume alcohol have a significantly worse prognosis than ALDH2*1 patients [43]. We therefore predict that ALDH2 dysfunction will accelerate HNSCC pathogenesis in habitual alcohol users through similar mechanisms related to this paper. Supporting this hypothesis, we have recently demonstrated that CSCs are enriched following alcohol exposure in HNSCC PDOs, although all samples were homozygous for ALDH2*1 [22]. Future studies will include ALDH2*2 HET samples. Finally, the ALDH2*2 polymorphism is associated with increased risk of colorectal cancer, gastric cancer, and breast cancer [44]. Although ethanol consumption has been demonstrated to enrich CSCs in these cancers, how ALDH2 status influences these phenomena is unclear [45].

There are several translationally relevant findings in this study. We provide proof-of-concept that pharmacological ALDH2 activation can reverse alcohol-induced tumor growth (Figure 4). Therefore, these activators may have efficacy as single agents. Of note, a next generation ALDH2 activator, FP-045, is currently in phase 2 clinical trials to treat patients with Fanconi Anemia (FA) (Trial Identifier: NCT04522375), a rare genetic disorder featuring bone marrow failure, leukemia, and young-onset HNSCC and ESCC (REF 48 here?). These inhibitors may also be used in combination with chemotherapeutic agents, given our findings that ALDH2 activation mediates sensitivity to cisplatin (Figure 5).

Furthermore, our findings may translate to other disease types. ALDH2 dysfunction exacerbates Fanconi Anemia and accelerates bone marrow failure through accumulation of aldehydic DNA damage [46,47]. We have previously demonstrated that the FA pathway repairs acetaldehyde-induced damage that may accumulate following alcohol exposure (Figure 3) [9]. Future studies will address whether the mechanisms delineated herein, including acetaldehyde-mediated CD44H cell enrichment, potentiate the increased risk for SCC pathogenesis in FA patients.

In addition to ALDH2, our data suggest that CD44 is an attractive therapeutic target. We demonstrate that CD44H cells survive high levels of cisplatin and may drive drug resistance. We therefore predict that targeting CD44H cells may improve patient outcomes. Several ongoing studies are addressing this possibility. A recent phase I trial (Identifier: NCT03078400) determined that 72% of patients with platinum-resistant ovarian cancer benefited from combined treatment with the CD44 inhibitor, SPL-108, and paclitaxel [48]. In addition to small molecule CD44 inhibitors, antibody-based therapies have also been tested in a variety of tumor types [49,50]. Furthermore, hyaluronic acid-based nanocarriers of cisplatin have been developed to specifically target CD44H cells [51]. Together, these ongoing efforts highlight the translational potential of our findings.

In conclusion, we demonstrate that a common SNP accelerates ESCC pathogenesis in response to a common risk factor of the disease. ALDH2 dysfunction promotes CSC enrichment and tumor growth following short term alcohol exposure. This phenomenon emerges in pre-malignant IEN, highlighting its importance during disease progression. Furthermore, ALDH2 dysfunction modulates response to a commonly used frontline chemotherapy, cisplatin. Both cisplatin and alcohol-induced CSC enrichment occur concomitantly with ROS accumulation, a well-established driver of this process. Treatment with a pharmacological activator of ALDH2 inhibits these deleterious phenotypes, demonstrating the therapeutic potential of our findings. Together, these data establish a link between ALDH2 dysfunction, CSC enrichment, and the progression of ESCC.

## MATERIALS AND METHODS

### Mice and murine preneoplastic esophageal organoids

C57BL/6 mice carrying Aldh2*1/Aldh2*1 (Aldh2 WT) and Aldh2*2/Aldh2*2 (Aldh2^E487K^ knock-in mice [52], gift of Drs. Che-Hong Chen and Daria Mochly-Rosen, Stanford University) were crossed to generate Aldh2*2/Aldh2*1 (Aldh2*2 HET) mice. Under IACUC-approved protocols, Aldh2 WT and Aldh2*2 HET mice were subjected to treatment with 100 mg/mL 4-nitroquinline-1-oxide (4NQO) (TCI NO250, Tokyo, Japan) in drinking water for 16 weeks to generate preneoplastic esophageal 3D organoids according to detailed protocols published by us [16,27]. In brief, 4NQO-treated mice were euthanized and dissected to collect esophagi and esophageal epithelial sheets. A portion of the tissue was reserved for histological diagnosis of intraepithelial neoplasia lesions. Following mechanical and enzymatic dissociation, the resulting single cells were suspended in advanced DMEM/F12 (ThermoFisher 12634028)-based Murine Organoid Media (MOM) [27]. 2×10^4^ single cells were embedded in Matrigel Basement Membrane Extract (Corning 354234) in a 24 well plate (ThermoFisher 12-556-006) and grown in 500 µL MOM media added into each well. Murine organoids were grown for 7-11 days with media changes every 48 hours and passaged or analyzed as described above for human ESCC cell line organoids.

### ESCC cell lines and organoid culture

All cell culture equipment and reagents were purchased from Thermo Fisher Scientific (Waltham, MA, USA) unless otherwise noted. ESCC cell lines TE11 (a gift of Dr. Tetsuro Nishihira, Tohoku University School of Medicine, Sendai, Miyagi, Japan) and TE14 (RCB2101; Cellosaurus Expasy CVCL_3336) (RIKEN BioResource Research Center Cell Engineering Division/Cell Bank, Tsukuba, Ibaraki, Japan) [53] were grown in monolayer as well as three-dimensional organoids utilizing RPMI-1640 supplemented with 10% fetal bovine serum as described previously [22,23]. TE11 and TE14 organoids were grown for 11 days with media changes every 48 hours. Organoid number was determined by capturing brightfield images using the Keyence Fluorescence Microscope BZ-X800 and Keyence software was utilized to determine the number and size of organoids, defined as spherical structures ≥5000 µm^2^ in size. Organoid formation rate (OFR) was determined as the percentage of the number of organoids formed at day 11per total number of cells seeded at day 0. Organoid morphology was determined via phase contrast imaging using the Evos FL Cell Imaging System. Mature organoids were passaged or collected for subsequent analyses as described [22,23,54].

Organoids were grown in the presence of the indicated concentration of EtOH and incubated for the indicated time periods. 100% EtOH (Decon Labs) was added directly into the media. To prevent evaporation of EtOH, plates were sealed with PARAFILM® M (Sigma-Aldrich). To control for any off-target effects of sealing plates, control wells (0% EtOH) were sealed as well. Organoids were treated with cisplatin (Santa Cruz sc-200896) for 72 hours.

### Immunohistochemistry and H&E staining

Organoids were treated as indicated, collected, centrifuged for 5-10 seconds at 2000 x g, and resuspended in 4% formaldehyde overnight at 4°C. Organoids were then embedded in 2% Bacto-agar (BD 214010) and 2.5% gelatin (ThermoFisher 35050061) prior to being processed for routine paraffin embedding and formalin fixation as described previously [17]. Tissues were subjected to paraffin embedding and formalin fixation immediately following harvesting. All samples were stained with hematoxylin and eosin as described [17]. For IHC, organoids were incubated with anti-Phospho-histone H2Ax (Ser139) (Cell Signaling 9718) at 1:250. All H&E and IHC data were evaluated by a board-certified pathologist.

### Flow Cytometry

Flow cytometric analysis of CD44 cell content was performed as described previously [22]. Briefly, organoids were subjected to the indicated treatment and dissociated into single cells. Cells were washed with PBS and resuspended in PBS containing 1% bovine serum albumin (Sigma-Aldrich A7906). TE11 and TE14 cells were stained with phycoerythrin-conjugated CD44 antibody (1:10, BD Biosciences 555479 Clone G44-26) in dark on ice. Murine cells were stained with an allophycocyanin-conjugated CD44 antibody (BD Pharmingen 559250). Dead cells were stained by DAPI (ThermoFisher) except in Annexin V assays, which used propidium iodide (Biolegend 640914). Data were analyzed using FlowJo software v10.7.1 (Tree Star). Untreated cells in the top 10% of intensity were defined as CD44H and cells in the bottom 10% of intensity were defined as CD44L cells. The CD44 gate was set using these values and applied to all experimental groups to determine CD44L and CD44H content following the indicated treatment.

Apoptosis was assessed via the FITC-conjugated Annexin V Apoptosis Detection Kit (ThermoFisher) according to manufacturer’s instructions.

### Mitochondrial Superoxide Assays

TE11 and TE14 cells were grown in monolayer culture as described [9], subjected to the indicated treatments, and then incubated with 5 µM MitoSOX™ (M36008; Thermo Fisher Scientific) and 20 µM Hoechst 33342 (ENZ-52401, Enzo Life Sciences) for 30 minutes at 37 °C in a humidified incubator. MitoSOX and Hoechst fluorescence intensity was assessed by imaging with a Keyence microscope and analyzed with ImageJ2 software (Version 2.9.0/1.53t). MitoSOX fluorescent intensity was background corrected and then normalized to the number of nuclei in the sample. Aldh2*1 and Aldh2*2 murine organoids were grown in 3D culture, subjected to the indicated treatments, collected, and dissociated into single cells. Single cells were incubated with MitoSOX as above and 4’,6-diamindino-2-phenylindole (DAPI) to detect dead cells. MitoSOX intensity was calculated as the average of fluorescent intensities in live cells.

### Xenograft Transplantation

Xenografts were performed as described [22] with the approval of the Institutional Animal Care and Use Committee at Columbia University. TE11-RFP and TE14-RFP cells were grown in monolayer culture, collected via trypsinization, and suspended in 50% Matrigel® Basement Membrane Matrix. The cell suspension was implanted subcutaneously into the dorsal flanks of 8-weel old athymic nu/nu mice (Taconic Biosciences, Hudson, NY, USA). 10% EtOH was included in the drinking water ad libitum from week 2 to week 6 following transplantation. Tumor volume was measured weekly with a digital caliper and calculated as follows: Tumor volume (mm^3^) = [width (mm)]^2^ x [length (mm)]^2^ x 0.5. During the EtOH treatment period, Alda-1 (20 mg/kg/day) or vehicle control (40% polyethylene glycol/10% dimethyl sulfoxide/ 50% water) was intraperitoneally injected two times per day. At week 6, tumors were dissociated into single cells and analyzed by flow cytometry for CD44.

### Statistical Analyses

Data presented in Figures 1-4 were analyzed/graphed with GraphPad Prism 8.0 software using the Student’s t-test. Data presented in Figure 5 were analyzed/graphed with the following: inkscape (1.2.1), matplotlib (3.4.3), numpy (1.22.4), python (3.8.8), pandas (1.4.3), scipy (1.9.0), seaborn (0.12.1), statsannotations (0.4.4).

## Figure Legends

**Supplemental Figure 1:**
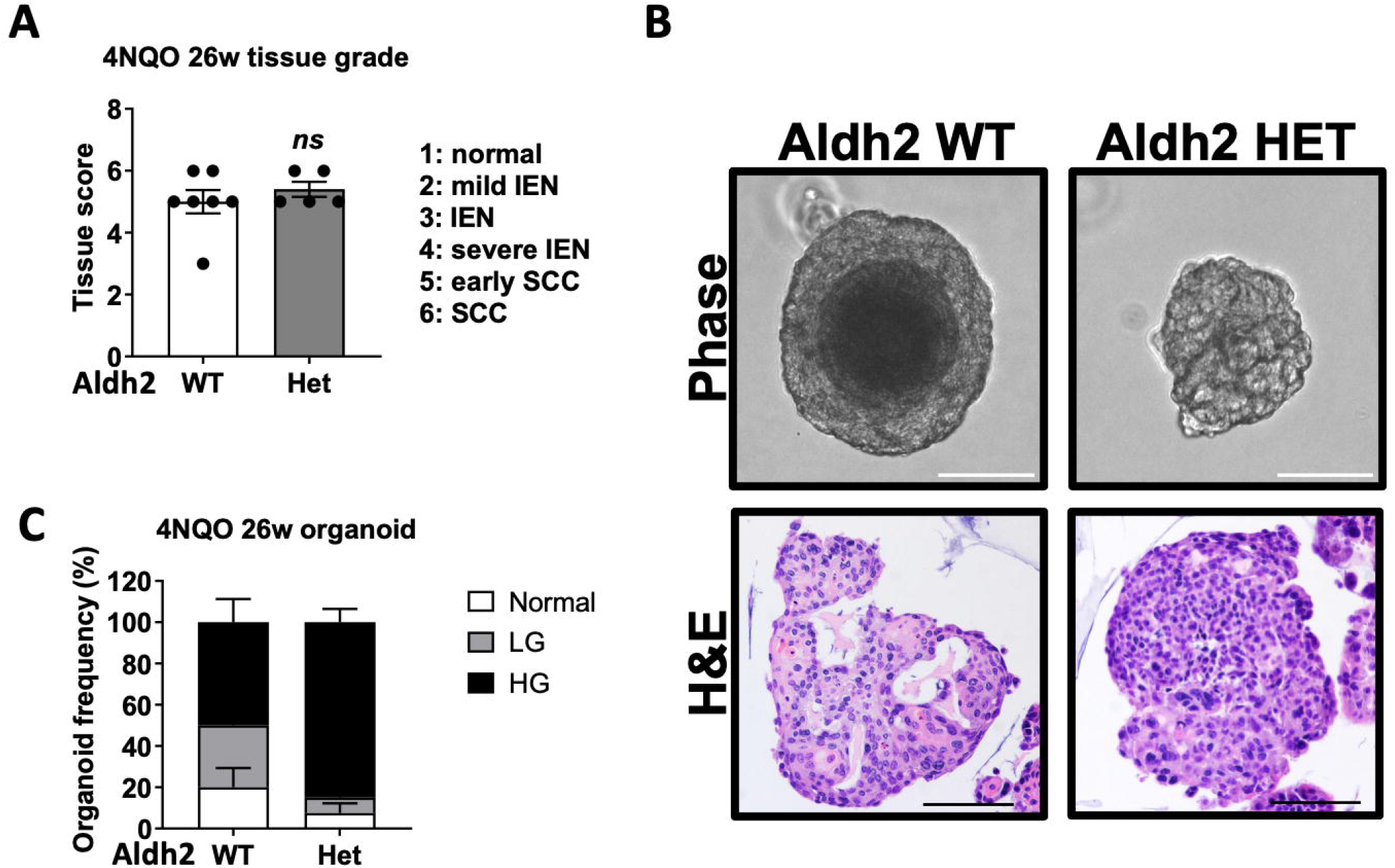
Establishing an isogenic organoid model of Aldh2 dysfunction during ESCC pathogenesis. (**A**) Mice were treated with 4NQO for 16 weeks following by an observation period of 10 weeks. Following completion of the experiment, esophagi were collected, stained by H&E, and diagnosed by a board-certified pathologist. The remaining tissue was used to make organoids (**B**) which were further H&E evaluated by the pathologist (**C**) Scale = 100 µm.

